# Pupil size reveals the perceptual quality and effortless nature of synesthesia

**DOI:** 10.1101/2025.11.24.690102

**Authors:** Christoph Strauch, Casper Leenaars, Romke Rouw

## Abstract

Synesthesia describes cross-over processes that can generate ’extra’ conscious percepts, such as seeing additional color when reading numbers. While existing research focuses on the mechanisms and effects of synesthetic associations, it often overlooks its most distinctive feature: unique sensory phenomenology. Here, we introduce pupillometry as an objective physiological measure of synesthetic color phenomenology. Across 16 grapheme-color synesthetes and two matched control groups, pupil responses tracked the brightness of synesthetic colors under constant physical visual input, scaling with self-reported strength. Synesthetic colors elicited pupil dynamics comparable to real colors, dissociating synesthetes from non-synesthetes. These responses emerged too rapidly to reflect imagery and scaled with reported color brightness, revealing cross-over caused genuine perceptual processing. Controls required to generate color associations showed greater effort-linked pupil dilation than synesthetes or controls who did not report colors, providing evidence for the effortless nature of synesthesia. Synesthesia thus provides a tractable human model for studying physiologically measurable phenomenology.

**Significance statement:** How can we measure what someone actually experiences? Synesthesia, in which stimuli such as numbers can evoke cross-over sensations like seeing colors, provides a rare test case. Here we make this extraordinary sensation objectively measurable and show that it has a distinct sensory signature. We find that pupil size reflects the brightness of synesthetic colors even when physical light remains constant. Pupils constrict for bright colors and dilate for dark ones, revealing the quality and strength of the percept. These responses emerge rapidly and differ from those of non-synesthetes. Together, synesthesia provides a tractable model for studying internally generated sensations, with pupillometry offering a direct, measurable window into conscious perception.

## 1 Main

Two observers presented with the exact same visual stimulus may experience a qualitatively very different percept [e.g., “the dress” 1]. While intuitively this may seem puzzling, theories on consciousness and vision explain such phenomena by noting that human perception, rather than a direct reflection of the external (shared) world, emerges in a constructive process of adjusting incoming sensory signals against our idiosyncratic expectations, knowledge and experiences [2–6]. This constructed understanding becomes especially apparent in color perception; even though color space is clearly defined in its physical and perceptual dimensions [7], it still fails to capture the “*what-it-is-like*” aspect of an individual’s color experience [1, 7]. Not surprisingly, being able to measure, understand, and eventually describe the emergence of these subjective qualia is therefore one of most hotly debated questions in the neurosciences, philosophy and beyond [5, 8–10]. Here, we present a natural experiment of subjective color phenomenology; a condition called grapheme-color synesthesia [11–14]. For grapheme-color synesthetes, certain linguistic inducers (e.g., grapheme ‘4’) automatically and consistently trigger additional and idiosyncratic conscious color percepts (e.g., a bright-blue color) alongside the veridical sensory input.

The different synesthesia types all share the defining characteristics of an additional conscious and consistent experience. Synesthetes can verbally report their additional experience, and synesthetic sensations can be measured in behavioral paradigms such as the ’synesthetic Stroop’ effect, or brain activation patterns in sensory cortex [15]. Furthermore, test-retest paradigms show how synesthetic, but not non-synesthetic associations are highly specific and consistent [16–18]. Thus, over the past decades, research has established synesthesia as a ’real’ condition that can reliably be identified using behavior, neurophysiology, and neuroimaging [11, 13, 15–19]. The most remarkable aspect of synesthesia is the subjective perceptual phenomenology of the induced additional sensation, i.e., color in grapheme-color synesthesia. This sets synesthetic sensations apart from (color) memory, thought, or amodal association. Synesthesia can thus offer an interesting doorway into examining qualia, the subjective perceptual phenomenology or first person (what’s-it-like) perspective. Furthermore, much like ordinary perception, the synesthetic experience is described as ’automatic’ in the sense that it comes effortlessly [13, 20, 21] [albeit not pre-attentive, see 22, 23]; the concurrent synesthetic color is ’just there’, even if incongruent with the task at hand [15, 20, 21]. Because each synesthete has a stable set of grapheme–color pairings, the color phenomenology can be examined independently of the physical properties of the inducing stimuli. Therefore, synesthesia might provide a unique window into how the brain’s constructive processes can generate additional, conscious content, cross-over experiences, often across modalities, going all the way down to the level of sensory phenomenology.

The measurement of such sensory phenomenology primarily relies on subjective reports and introspection, methods often criticized for potential unreliability and susceptibility to biases or expectancy effects [e.g., 24–26]. These concerns extend to synesthesia research, where objective measurements are called for to corroborate subjective reports [27–29]. Instead, current paradigms capturing synesthesia employ objective measures, but fail to capture its phenomenology [16, 21, 22, 30]. Behavioral and neurophysiological findings suggesting synesthetic colors behave like printed colors in turn have been questioned regarding replicability and interpretability [23, 31–33]. In short, the lack of agreed-upon objective methodology is a critical roadblock obstructing scientific examination of the extraordinary synesthetic phenomenology. By extension, this locks the condition’s potential to inform our understanding of the constructive top-down cross-over processes that can generate additional conscious percepts.

We propose that pupillometry is the tool to break this gridlock. Pupil size constricts in response to externally increased brightness and dilates when brightness decreases. Remarkably, akin to our color phenomenology not directly following from physical color input, the pupillary light response does not strictly follow the physical light entering the eye, but reflects the percept as interpreted by the viewer [34–42]. Research has revealed such sensitivity to perceptual phenomenology in unimodal contexts [e.g., phenomenological vividness of an (imagined) visual image 40, 43, 44]. Building on this evidence, we hypothesized that the cross-over color phenomenology in synesthesia, if truly sensory in nature, could likewise be inferred from changes in pupil size. Hereby, the direction and magnitude of these changes should provide a scaled response reflecting the brightness of the experienced synesthetic colors; pupillometry may thus provide both qualitative and quantitative characterizations of the synesthetic color phenomenology (see Figure 1a and b for an illustration of the rationale and proposed mechanism). If synesthetic cross-activations indeed reach all the way down to low-level (sensory) processes, pupillometry can track their precise temporal onset, as well as provide scaled measurements of their phenomenological properties (i.e., the relative change in pupil size corresponding to the brightness of the synesthetic color).

**Figure 1:**
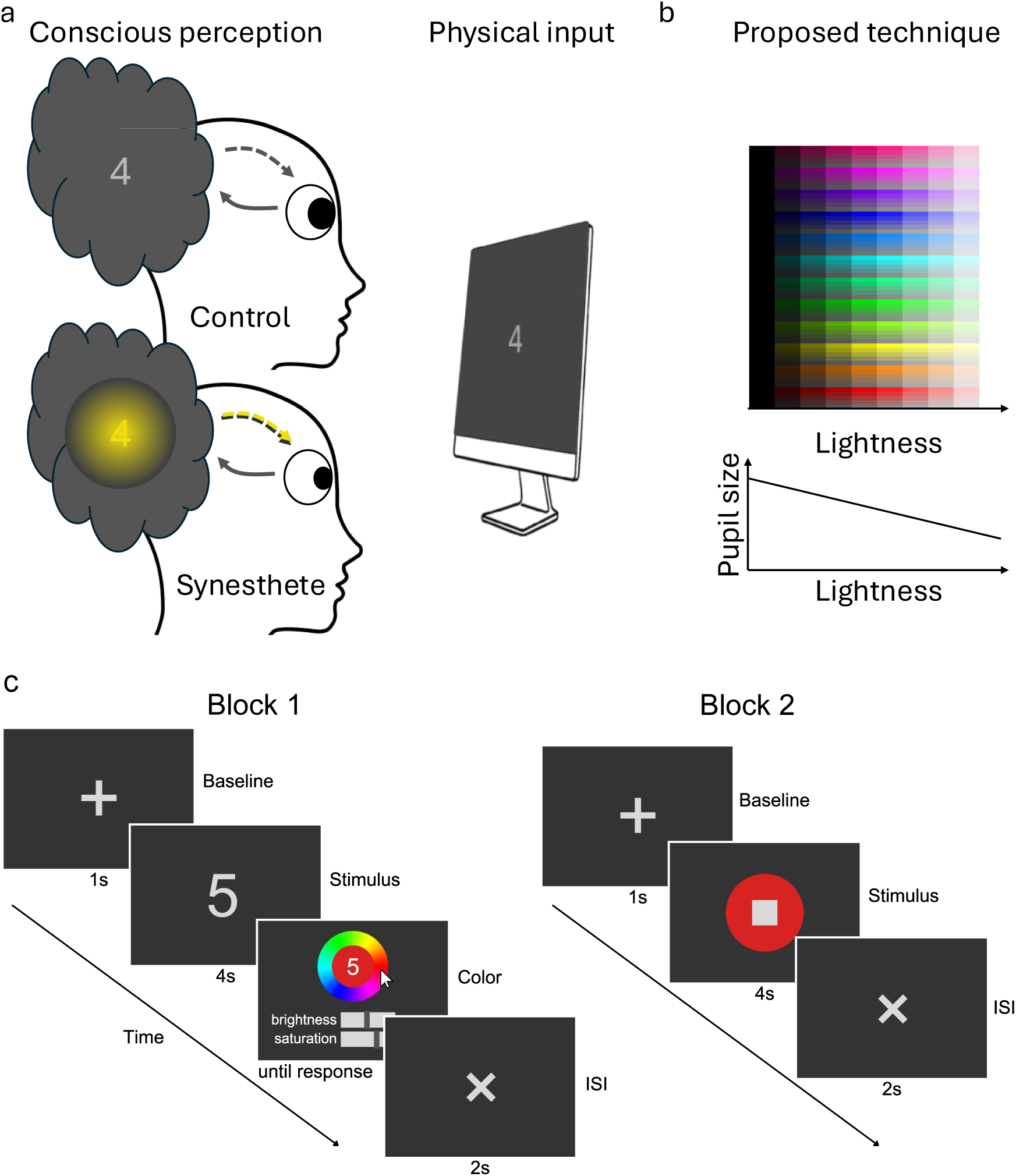
Mechanism and paradigm. **a** phenomenology results from external (solid arrow) and internal contributions (dashed arrow). The integrated brightness should affect pupil size: Light (dark) synesthetic colors should cause constrictions (dilations) at equal physical luminance in synesthetes, but not in controls where externally and internally generated brightnesses align. **b** We expected synesthetes’ pupils to be larger for reported lower brightness and smaller for reported higher brightness. **c** Paradigm. Block 1: a digit was presented. Participants (except passive controls) subsequently indicated the color that most closely corresponded to the digit in their opinion. This was followed by an interstimulus interval (ISI). Block 2 (synesthetes only): a disk was presented, colored according to the synesthete’s average indicated color for that digit. At its center sat a gray patch matching the luminance and pixel area of the original digit from Block 1, together allowing assessment of externally triggered light responses.

To investigate this, we tested 16 grapheme-color synesthetes and two control groups of 16 participants each. Participants viewed graphemes (digits) on a computer screen during eye-tracking, and indicated, after each trial, which color most closely matched with the respective grapheme (see Figure 1c for paradigm).

## 2 Results

### 2.1 More consistent and strongly coupled colors in synesthetes

Selected colors are visualized per participant in Figure 2a. Synesthetes reported colors more consistently (*t*(30) = 9.910, *p* < 0.001, *d* = 3.504, 95% CI = [2.370, 4.614] (in line with previous work, see [see e.g. 16, 45, 46]) and more strongly coupled to graphemes (*t*(30) = 12.690, *p* < 0.001, *d* = 4.487, 95% CI = [3.150, 5.801]) than controls (see Figure 2b). There was considerable variation in reported color lightness for all participants (see Figure 2c, d), setting the basis for a possible inference of color phenomenology via the pupil light responses in both synesthetes and controls. Importantly, reported lightness of colors almost exactly matched between controls (*M* = 0.478, *SD* = 0.065, lightness scaled between 0 = black and 1 = white) and synesthetes (*M* = 0.479, *SD* = 0.070; *t*(30) = 0.031, *p* = 0.975, *d* = 0.011, 95% CI = [-0.704, 0.682]; *BF*_01_ = 2.973). Together, our synesthete participants were grapheme-color synesthetes as per the gold standard of the field [16], reporting specific, strong and consistent colors in response to graphemes.

**Figure 2:**
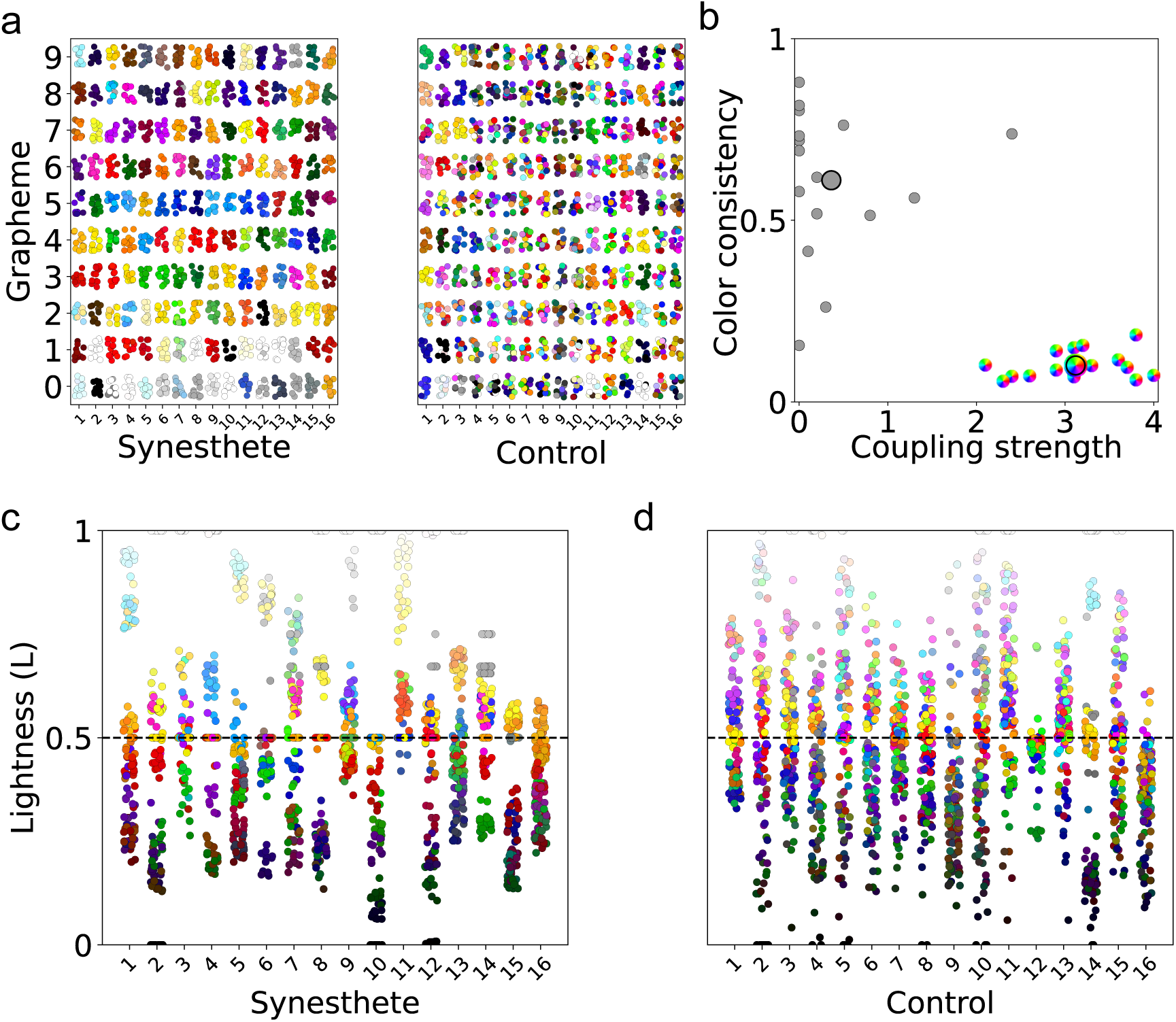
**a** Reported colors per grapheme on all trials for synesthetes (left) and controls (right). **b** Synesthetes showed (near) perfect grapheme-color consistency and moderate to very strong grapheme-color couplings (rainbow circles), while controls reported none to moderate coupling and varied in consistency (grey circles). Note that higher consistency is reflected in lower color distance, hence lower values [17]. Larger dots indicate group means. **c,d** (HS)Lightness of color reports per synesthete (c) and control (d). Blacked dashed line represents lightness being 0.5. See Supplementary Figure 1 for color reports on the hue and saturation axes.

### 2.2 Pupil responses reveal the quality of synesthetic color perception

Having established similar reported-color lightness levels between active controls and synesthetes, we next investigated whether the pupil light response betrays synesthetic color. Perceived, (covertly) attended, or even imagined brightness modulates pupil size in the same direction as changes in physical luminance - i.e., in both directions, constriction *and* dilation [see 47, 48, for reviews]. We therefore expected bright perceived synesthetic colors to be betrayed by relative pupil constriction and dark synesthetic colors to be betrayed by relative pupil dilation. We did not expect non-synesthetes to show pupil size alterations in accordance with the brightness of their associated colors. Pupil responses to reported color lightness were analyzed separately for synesthetes and active controls. A visual inspection of per-timepoint demeaned pupil traces for participants having at least 25 trials in above and below median lightness bins respectively (Figure 3) demonstrates larger pupil sizes for dark graphemes and smaller pupil sizes for light graphemes in synesthetes (mid row, Block 1). In controls, this was very weak, if present at all (top row, Block 1). As expected, when splitting pupil size for colored discs similarly along lightness, synesthete pupil size demonstrated descriptively even larger and earlier changes than for synesthetic color. Dependent-samples t-tests for averaged pupil size in response to graphemes (stimulus interval) between 800 ms and 4000 ms, split by (reported) color lightness, showed different pupil responses for synesthetes both for synesthetic colors (Figure 3d, *t*(11) = 4.669, *p* = 0.001) and externally triggered light responses in synesthetes (Block 2, see Figure 1c; Figure 3f, *t*(10) = 4.550, *p* = 0.001), but not in controls (Figure 3b, *t*(12) = 0.850, *p* = 0.412). Note that the effects here are visualized as counterfactuals. So while the pupil dilated for dark relative to bright experienced colors in synesthetes, this does not mean that the pupil net dilates and constricts to dark and bright experienced colors relative to baseline, but only relative to the counterfactual (see Supplementary Figure 4 for net pupil size changes).

**Figure 3:**
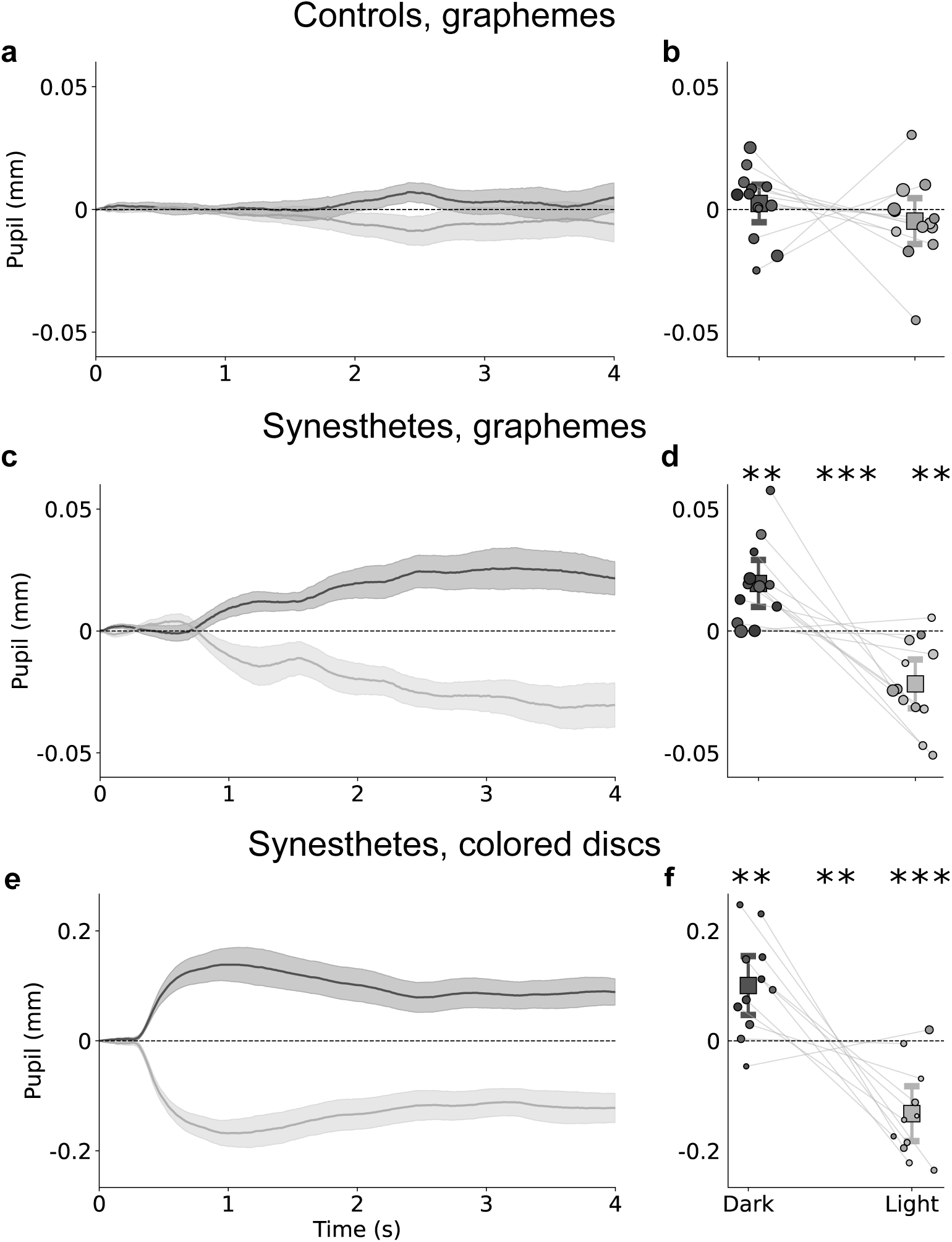
Pupil size change to graphemes, median-split by reported color lightness (dark gray = low lightness; light gray = high lightness). Top row: pupil responses to graphemes in controls. Mid row: pupil responses to graphemes in synesthetes. Bottom row: pupil responses to colored discs in synesthetes (Block 2). **a, c, e** Depict average, baseline-corrected and within-participant demeaned pupil responses. Shaded error bands: ±1 SE across participant means. **b, d, f** depict mean pupil size (800–4000 ms) for dark vs. bright colors. Dots show individual participants; squares denote grand means with 95% CIs as whiskers. Dot luminance corresponds to the participant’s average synesthetic color lightness per bin, dot size to the number of trials. **: p < .01, ***: p < 0.001 based on within samples and one sample t-tests. Significance relative to zero for lightness bins (left, right) and between bins (center). Participants with less than 25 trials per bin excluded for visualization (controls: *n* = 3, synesthetes: *n* = 4, see **a, c** and Supplementary Figure 4 for visualization without demeaning and Supplementary Figure 3 f^9^or visualization without data exclusion).

#### 2.2.1 Pupil size betrays the lightness of synesthetic color

To optimally account for the data structure, we next ran a linear-mixed effects model (LME) predicting pupil size. The LME effectively considers all trials and was fitted for the average pupil size between 800 ms and 4000 ms (reported in brackets) as well as separately for every timepoint in the stimulus interval (visualized in Figure 4). Starting from a full model containing all interactions between the following factors, the final LME for synesthetes (for both discretized and per-timepoint analyses) was determined using AIC-based backward selection while retaining coupling strength (self-reported strength of color-grapheme coupling) and PA score (projector-associator score, Rouw & Scholte [49]) as predictors for synesthetes, predicting average pupil size between 800 ms and 4000 ms (Wilkinson notation: *Pupil size* ∼ *grapheme + lightness + coupling strength + PA score + lightness * coupling strength + lightness * PA score + lightness * coupling strength * PA score + (1 | participant)*). The effects of all predictors over time on pupil size are depicted in Figure 4a (Supplementary Figure 6 for controls). We found significant modulations of pupil size by the lightness of the grapheme’s synesthetic color - sustained and in the to-be-expected time window. Specifically, the pupil constricted more for brighter reported colors, and dilated more for darker reported colors, as predicted (Average pupil size 800-4000 ms, *t* = -3.601, *p* < 0.001). In an LME ran for synesthetes and controls and using only graphemes and lightness as predictors, we found lightness to predict pupil size in synesthetes (*t* = -2.844, *p* = 0.004), but not controls (*t* = -0.606, *p* = 0.544). However, when taking group as interacting factor in a joint LME, there was no interaction of lightness and group (*t* = -0.949, *p* = 0.342). Together, this demonstrates that 1) the pupil reveals the hidden qualia of synesthetic color (along the brightness axis) and 2) that such perception recruits the very same networks as are active during the perception of ’real’ differently luminant stimuli itself, down to such a level that even the sensory organ is affected.

**Figure 4:**
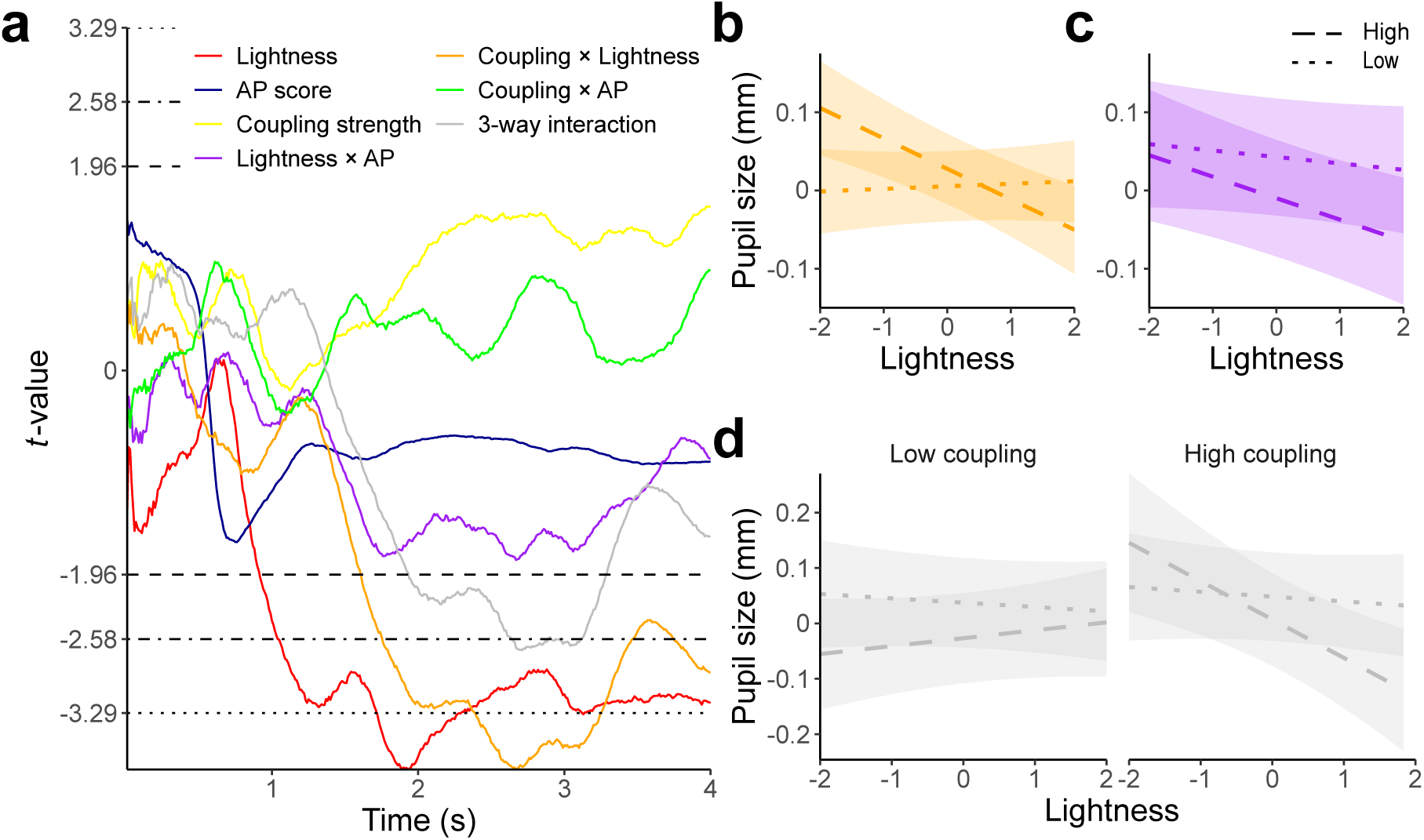
Results of per-time-point linear mixed effects model (LME) predicting pupil size in synesthetes while presented with graphemes. Covariates for the individual graphemes and intercept are not visualized here. **a** depicts t-values of the LME over time. Horizontal lines denote significance threshold (*p* = 0.05 dashed, *p* = 0.01 dot-dashed, *p* = 0.001 dotted). Higher lightness was associated with smaller pupil size (red), this effect was stronger for stronger reported grapheme-color couplings (orange), with a trend for higher PA scores (purple). Furthermore, higher lightness constricted the pupil more for stronger grapheme-color couplings in synesthetes with higher PA scores (gray, three-way interaction). **b-d** visualize interactions for the LME run on the average pupil size between 800 ms and 4000 ms. Dotted denotes low, dashed high of median splits. **b** Interaction of grapheme-color coupling strength with lightness: lightness affected the pupil more when grapheme color couplings were reported higher. **c** Interaction of PA scores with lightness: lightness affected the pupil more for synesthetes with higher PA scores, but note that this effect only reached borderline significance for a short interval. **d** three-way interaction of lightness, coupling strength, and PA score. See Supplementary Figure 6 for the same model in controls.

#### 2.2.2 Lightness of stronger color-grapheme couplings affects pupil size stronger in synesthetes

The pupil response to synesthetic color lightness was amplified for stronger reported coupling strengths between graphemes and colors (interaction coupling strength * lightness: per-timepoint from ±1700 ms, see Figure 4b); average pupil size 800-4000 ms: *t* = -3.093, *p* = 0.002). In other words, synesthetes have metacognitive insight into the 3) *strength* of their synesthetic color perception as revealed by the pupil response. While a trend for stronger effects of lightness for more projecting synesthetes was observed (average pupil size 800-4000 ms: *t* = -1.421, *p* = 0.155, see Figure 4c), that might mean that the nature of color percepts affects the pupil response to synesthetic color lightness, our data cannot answer this question conclusively. Finally, effects of reported color lightness on pupil size were stronger with higher indicated grapheme-color couplings for individuals with higher PA scores, i.e., less associating, more projecting synesthetes (three-way interaction, ±1900 ms-3300 ms; but not for the average pupil size 800-4000 ms: *t* = -1.779, *p* = 0.075, see Figure 4d). Together, the pupil therefore revealed both *quality and quantity* of self-reported synesthetic colors.

For controls a separate model was run, now without the PA score as predictor (not assessed for controls). Neither lightness (*t* = -0.815, *p* = 0.415), coupling strength (*t* = 0.438, *p* = 0.661), nor their interaction gained significance (*t* = -1.058, *p* = 0.290; all for average pupil size between 800 ms and 4000 ms). Critically, we also ran a LME with the three-way interaction of coupling strength, group, and lightness (Wilkinson notation: *pupil = grapheme + group + lightness * group + coupling strength * light-ness * group + (1 | participant)*). This analysis revealed a significant three-way interaction between lightness, coupling strength, and group (*F* = 3.86, *p* = .021), indicating that the lightness × coupling strength effect on pupil size was not equivalent across groups. Decomposing this interaction by group, the lightness × coupling strength slope was significant in synesthetes (*t* = -2.59, *p* = .010) but not in controls (*t* = -1.01, *p* = .311), suggesting that reported lightness and its coupling strength were more consistently related to pupil size in synesthetes than in controls. Note however, that this decomposition does not directly test whether the two slopes significantly differ from each other. We found pupil size to be marginally larger in controls than in synesthetes (*t* = 1.94, *p* = .062; see later sections for more in-depth analyses).

Lastly, we tested whether higher color consistency, the gold-standard assessment of synesthesia [16], predicted stronger pupil responses according to color lightness. It did neither in synesthetes (significant predictors: lightness and interaction of coupling strength and lightness; very limited variance of consistency) nor in controls (no other significant predictors; see Supplementary Material for full analyses). Together, this demonstrates that the consistency of colors was, in this study, not found related to the pupil responses to color brightness.

### 2.3 Pupil responses demonstrate the automaticity of synesthetic colors

Having established the qualia and quantity of synesthetic colors through pupillary responses, we next turned to the other core defining feature of synesthesia: its presumed *effortless* nature. Synesthetes report the colors to emerge automatically - rather than via active and effortful cognitive (e.g., memory) processes.

#### 2.3.1 Synesthetic colors affect pupil size delayed relative to physically presented colors

In Block 1, synesthetes viewed individual digits, and as the pupil revealed, an evoked characteristic (synesthetic) color. In Block 2, we then physically presented, for each digit, its previously determined average synesthetic color as a colored disc on the screen. At the center of each disc sat a gray rectangle whose luminance and pixel area matched the original digit’s lightness and size from Block 1 (see Figure 1c). As expected, physically presented color discs let the pupil constrict strongly in response to bright and dilate in response to dark colors, respectively. Equally expected, this effect was numerically and statistically more pronounced than the response to synesthetic colors [akin to stronger effects for direct fixation compared with covert attention only, see 37]. Interestingly, the time course of the pupil response to physical vs synesthetic color differed markedly. Specifically, pupil size first responded significantly to physical luminance after 330 ms (see Supplementary Figure 7 for per-timepoint LME; in line with response latencies of similar control populations, see Koevoet *et al.* [39], Bergamin & Kardon [50], and Strauch *et al.* [51]), but only responded significantly to synesthetic lightness at about 870 ms (see also Figure 3c vs e and Figure 4 for per-timepoint LME). Assuming that internally generated lightness does elicit a pupil light response with similar latency as the physically triggered lightness reflex arc (330 ms here), this implies that synesthetic perception has to emerge within 540 ms, including the recognition of the digit itself. This fast emergence makes it highly unlikely that synesthetes imagined a color after processing a grapheme, as this must take up more time [52].

#### 2.3.2 Reporting colors to graphemes is more effortful for controls than for synesthetes

Lastly, we reasoned that synesthetic color perception should be relatively low in effort. We therefore exploited another, distinct feature of pupillary responses: In absence of any luminance changes, pupils *dilate* more from baseline the more effort is exerted [e.g. 48, 53–57]. Figure 5a visualizes pupil size change to baseline. Figure 5b depicts average pupil size change to baseline between 800 ms and 4000 ms per participant. Mental effort presents in task-evoked pupil dilations, yet other factors simultaneously affect the pupil, such as luminance and contrast changes at trial onset, as well as slower trends across the session (e.g., fatigue). To reduce the influence of these slower, non-trial-locked fluctuations while retaining the trial-evoked dynamics, we calculated the first derivative of the pupil time course to assess the velocity of pupillary changes (Butterworth filter, 18 Hz, order 3, 2.5 Hz lowpass, following our previous works [58, 59]). Figure 5c depicts this derivative, Figure 5d the average derivative per participant between 700 ms and 2000 ms (where we observed the effect). We found a stronger pupil dilation rate for active (reporting colors) compared with passive controls (not reporting colors; *t* = -4.254, *p* < 0.001) and for active controls compared with synesthetes (*t* = -2.828, *p* = 0.007), but no difference between synesthetes and passive controls (*t* = 1.424, *p* = 0.161, all on the interval 700 ms-2000 ms, LME formula in Wilkinson notation: *pupil = group + (1 | participant*)). Together, this demonstrates that having to report a color after seeing the grapheme is associated with pupil dilation in controls. This pupil dilation is absent in synesthetes and in controls not having to indicate any color. We interpret this effect to reflect differences in effort, not least because reported lightness was similar for synesthetes and active controls overall (see Figure 2c,d). This higher degree of effort in active controls as compared with passive controls and synesthetes may not be so surprising, given that the task to report a color when not seeing a color is not trivial. We argue that the obtained difference between active controls and synesthetes performing the exact same task is an additional consequence of the reported ’automatic’ (effortless) nature with which synesthetes can indicate colors for graphemes. Together, the time course of effects and the reduced degree of effort needed for synesthetes during the task provides converging (and perhaps conclusive) evidence that synesthetic percepts are indeed fast and effortless, in line with synesthetes’ subjective reports [46].

**Figure 5:**
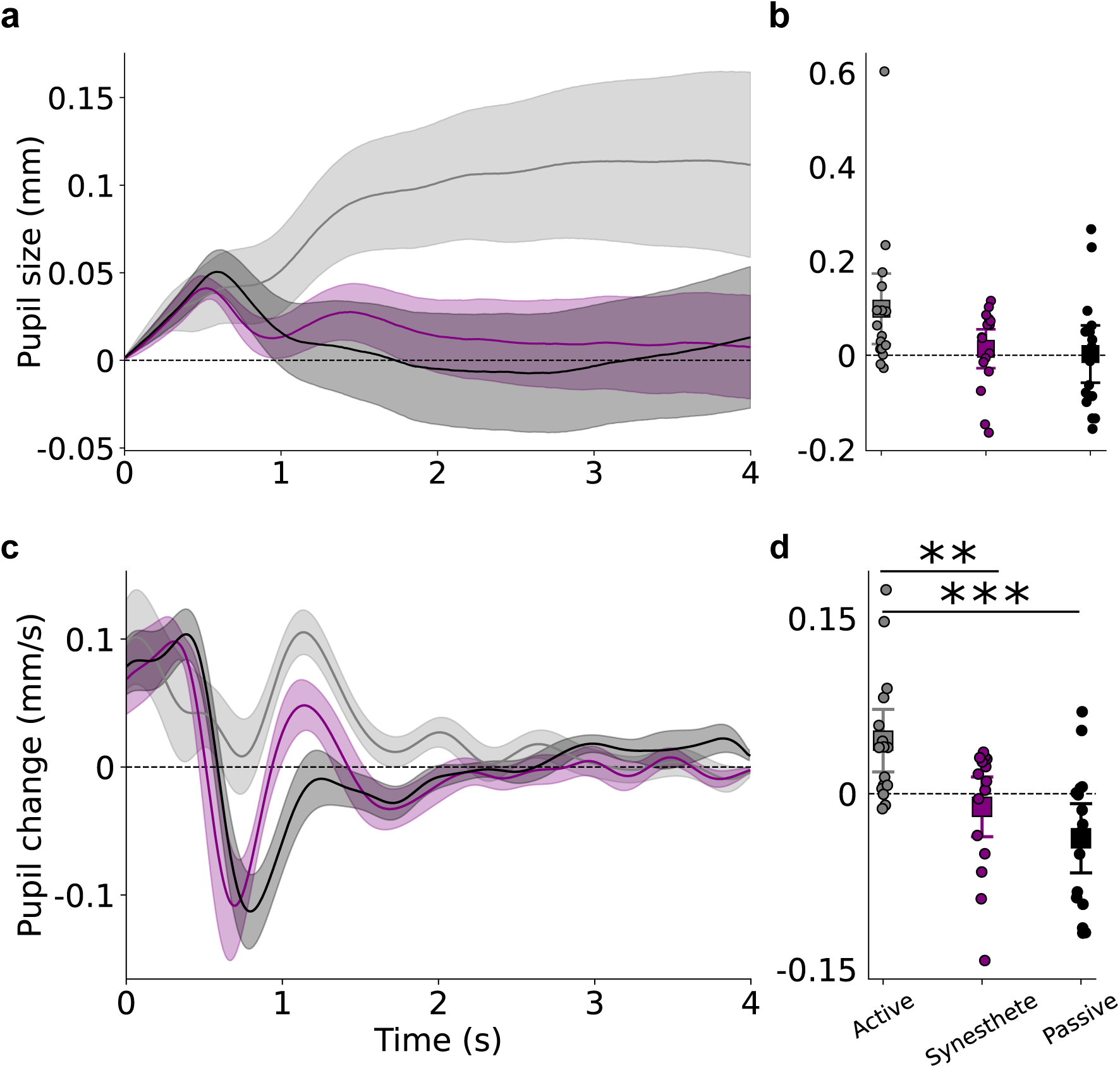
Average pupil responses to graphemes from baseline, split by group: controls picking a color forced-choice (’active’, gray), controls passively viewing the graphemes (’passive’, black), and synesthetes (purple). **a** Pupils dilated more for active controls than both synesthetes and passive controls. Shaded error bands represent 95% confidence intervals across participant means. Horizontal black line represents average pupil size during baseline. **b** Mean pupil size (0.8 s–4 s interval) per group and participant. Dots show individual participants; squares denote grand means with 95% CIs. **c** as a, but for the velocity of pupil size changes (first derivative, filtered). **d** as b, but for the velocity of pupil size changes and the 0.7 s-2 s interval. *p* < .01: **, *p* < .001: ***, based on two-sided independent sample t-tests.

## 3 Discussion

We demonstrate that pupil size changes reveal the qualia of synesthetic (grapheme-color) percepts. Specifically, pupils constricted when viewing digits that evoked brighter synesthetic colors and dilated to digits that evoked darker synesthetic colors. In contrast, non-synesthetes presented with the exact same physical input did not show modulation of pupil size to the brightness of their associated colors. Synesthetes (but not controls) showed high color consistency, in line with the diagnostic ’gold standard’ maintained in the field [16, 17]. While such standardized objective diagnostics [e.g., 15–17, 60] reliably separate synesthetes from non-synesthetes, our findings directly corroborate the most discerning and -arguably- most fascinating characteristic of grapheme-color synesthesia; the reported phenomenology of the synesthetic color.

This offers practical and theoretical progress in clarifying the boundary between synesthetes and non-synesthetes [61–63]. Conceptual cross-over correspondences, which are consistent at the group level, can also be observed in the general population [14, 64, 65]. Moreover, synesthesia-like Stroop effects can be induced in non-synesthetes through training [66, 67]. However, as illustrated by the traditional Stroop effect [68], neither consistent color associations nor Stroop-like conflicts depend on sensory color phenomenology. Our technique, linking synesthetic brightness and pupil size for the first time, maps out phenomenological features of cross-over (grapheme-to-color) experiences. In synesthetes, pupillometry tracked the qualia of associated colors along the brightness axis indicating that similar networks are engaged as during perception of ’real’ (printed) differently luminant stimuli.

The effect of color lightness on pupil size scaled with the indicated *strength* of individual grapheme–color couplings, physiologically validating synesthetes’ meta-cognitive insight into their own associations [69]. In future work, per-trial ratings could take this a step further by assessing moment-to-moment fluctuations and their neural correlates.

Along with the conscious perception of color, a main feature of synesthetic color experiences is that they happen automatically [e.g., 70]. Note that ’automaticity’ in this study means the reported effortless nature of the additional sensations, as synesthetic sensations are unlikely evoked pre-attentively (see [71]). As increases in mental effort are tightly coupled with pupillary dilations [e.g. 48, 53, 54, 56, 72], we could put this to a direct test. Indeed, we found faster pupil dilation in active controls than in passive controls at constant physical stimulation. This stronger effort-linked increase in pupil dilation in active controls (preparing to report a color after viewing the digit) as compared with passive controls (observing the same digits but without color task) most likely reflected the non-trivial nature of the color task for non-synesthetes. Furthermore, active controls exhibited greater pupil dilation and thus effort than synesthetes, even though both groups received the same color-reporting task. Pupil size change rates showed no significant difference between synesthetes (asked to pick a color after viewing digits) and passive controls (no subsequent task). Unlike non-synesthetes, synesthetes thus measurably experience their synesthetic colors effortlessly, as their conscious color phenomenology allowed them to see and pick the right color, much like non-synesthetes view an actual (typeface) color.

Effort-evoked pupil dilation also speaks to an important theoretical question, as synesthetes often report richer mental imagery [73, 74]; could our findings reflect an active color-imagery strategy [akin to 40, 43, 44] rather than automatic color emergence? We deem this highly unlikely. First, generating and maintaining mental images is effortful and produces according pupil dilations [43, 44, 53, 75, 76]. Second, the response timeline supports automatic generation. Synesthetic color affected pupil size after 870 ms. Assuming a constant pupil light response latency to external colors (here 330 ms, see [50]) and internally generated colors, plus at least 150 ms for digit recognition [77, 78], this leaves only 390 ms for imagery - which in turn has been shown to be slower [500 ms+, based on MEG, 52]. Our fast pupil responses support previous physiological studies showing early emergence of the synesthetic response [19], and a proposed synesthetic physiological mechanism of a recurrent loop from grapheme recognition to color perception during the forward sweep of visually presented information [79]. Future work could further specify this mechanism using pupillometry.

Pupillometry thus allows to measure several key aspects of synesthesia, most prominently the unique phenomenology, the precise temporal onset, time course and strength of these effects, but also the degree of effort evoked by color reports. The measure is further unobtrusive, relatively inexpensive, and has a high signal-to-noise ratio for a physiological marker. Perhaps most importantly, pupillometry physiologically tracks synesthetic brightness in the sensory organ itself. Compared to neuroimaging studies [12, 15, 49], pupillometry may offer a more direct window into synesthetic phenomenology, as the directionality between pupil light reflex and perceived brightness is straightforward. Finally, improved understanding of the underlying processes can be obtained by contrasting responses to perceived versus actual (physical) brightness, given that the pupil light reflex is a well-characterised reflex arc involving few inferential steps. Furthermore, this method adds a metric allowing between-trial or between-participant quantitative comparisons that is not present in subjective reports: if two individuals indicate seeing a ’bright-blue’ experience on a questionnaire, does this mean that their experiences are the same? Can their intensities ever be compared? Pupillometry thus extends the synesthesia research toolbox by providing an objective metric with shared measurement unit for the quality and strength of inducer-to-concurrent links, suited both across and within individuals. Future studies could examine to what degree training a non-synesthete to associate specific colors to particular inducers (e.g., digits), can provide similar patterns of results as genuine synesthesia [67, 80, 81]. Could learning produce similar brightness-related pupil effects in non-synesthetes? Similarly, would effort-linked responses diminish with increased training duration? The perhaps most interesting question relates to response latencies: Would a trained participant ever be able to produce brightness-related pupil effects as fast as a synesthete? Finally, pupil light responses in Block 2 were only assessed in synesthetes. While these closely match such of control populations [50, 82], subtle between-group differences cannot be excluded and could ideally be assessed in future and replication work.

Taking a broader perspective, this work further informs about cross-over processes: associations stored in a synesthete’s brain can guide the constructive mechanisms that generate additional conscious sensations. Consistent with this view, recent work has identified cross-modal predictive-coding mechanisms, in which unimodal predictions are integrated across distributed networks to form cross-modal representations [83]. We suggest that synesthesia may exemplify such a (often) cross-modal constructive process, extending to the level of additional sensory color phenomenology. Indeed, synesthesia is an important phenomenon in context of interdisciplinary consciousness research. Fascinatingly, a synesthete may see a presented number 4 as bright-blue, while knowing and seeing it as dark gray *at the same time* [84, 85]. Such idiosyncratic “extra” challenge the leading frameworks of consciousness. Integrated Information Theory [10] posits that each unified conscious moment corresponds to a single maximally integrated causal complex, yet (stronger projecting) synesthetes experience an additional color dimension bound to the same grapheme. Similarly, Global Workspace Theory holds that only one content is broadcast at a time [e.g. 9, 86, 87], yet shape and synesthetic color co-occupy the global workspace without interference. By varying the luminance of inducers while measuring the pupil light response in synesthetes, future work may disambiguate the (relative) strength of these two contributors to perception. Similarly, the here demonstrated perceptual nature of synesthetic perception may inform research into internally generated percepts and their properties. With the findings and technique proposed here, it remains inaccessible what it ’feels like to be a bat’ [88], yet we may come closer to objectively measure the seamless ’controlled hallucinations’ [5] that we experience as reality.

## 4 Methods

### 4.1 Participants, inclusion, and ethics

Sixteen synesthetes (*M*_age_ = 23.94, *SD*_age_ = 3.40, 13 women, 3 men), *n* = 16 age-matched control participants watching graphemes passively (*M*_age_ = 23.50, *SD*_age_ = 2.11, 9 women, 7 men), as well as *n* = 16 age-matched ’active’ control participants watching graphemes and indicating colors forced-choice (*M*_age_ = 24.75, *SD*_age_ = 4.20, 14 women, 2 men), all with otherwise normal or corrected-to-normal vision took part in the tasks. No participants were excluded. All participants had normal or corrected-to-normal vision without eye diseases. Synesthetes and controls were recruited through a snowball sample using a survey that was sent to thousands of people in the Netherlands and neighboring countries, using messenger apps and forums (Whatsapp, Signal, Reddit). Furthermore, the survey was distributed to a worldwide synesthesia network [89, 90] and participants with synesthesia of a concurrently running study at the University of Amsterdam were approached. Respondents indicated their form of synesthesia if present. Controls indicated not having synesthesia. The experimental procedure was approved by Utrecht University’s Faculty of Social Sciences ethical review board (24-0521). We herein predicted our main finding - the pupil light response to reflect the quality of synesthetic color. All participants gave written informed consent prior to participation.

### 4.2 Apparatus

Gaze position and pupil size were recorded at 1000 Hz using an Eyelink 1000 desktop mount (SR Research, Ontario, Canada) in a brightness- and sound-attenuated, mostly dark laboratory. A chin- and forehead-rest limited head movements. Stimuli were presented using PsychoPy [v.2024.2.3; 91] on an ASUS ROG PG278Q monitor (2560 x 1440, 100 Hz) positioned 67.5 cm away from eye position. The monitor was not linearized. The eye-tracker was calibrated and validated (7 points) at the beginning of the session and recalibrated whenever necessary throughout the experiment.

### 4.3 Procedure, task, and stimuli

Before the experiment, grapheme-color synesthetes indicated where they see their synesthetic colors, (in the mind versus in the outside world) on the ’projector-associator’ (PA) questionnaire, answering twelve questions on a five-point Likert scale [49]. Furthermore, participants (active controls and synesthetes) assessed for each grapheme separately the subjective strength of color-grapheme couplings on a 5-point scale from 0 (’None’, no color coupling) to 4 (’Very strong’, very strong color coupling). This rating is referred to as coupling strength in this manuscript. See Supplementary Materials for the questionnaire used.

The experiment started with calibration and validation of the eye-tracker. Next, participants began with Block 1 of the task (see Figure 1c). Participants were first presented with a fixation cross for 1 s on a dark gray screen, if gaze was successfully kept central during this baseline screen (within 1.5 °visual angle from the center), this was followed by a random single digit number (0-9, letter height: 1.42° visual angle), presented for 4 s. Digits occupying more physical space on the screen (e.g., ’8’) were presented less bright than digits occupying less space on the screen (e.g., ’1’), scaled between 65% and 75% of the luminance range of the screen. Subsequent to this decisive measurement phase, participants saw the same digit as before as a reference on the screen. Participants were asked to use their mouse to indicate hue, saturation, and lightness (HSL) using sliders. These sliders changed the color of a circle surrounding the reference digit. The herewith obtained per-trial lightness values (L of HSL) form the main predictor in the manuscript. Only synesthetes were allowed to press ’space’ to indicate the absence of any synesthetic color in which case the lightness of the screen background was used. Active controls were forced to always indicate a color, passive controls did not indicate a color. Participants were instructed not to blink or look away from the central character during baseline and stimulus phase to prevent pupil foreshortening errors and luminance confounds [48]. Consequently, trials containing blinks or gaze deviating more than 1.5°visual angle from the center were discarded and had to be repeated in random sequence until 120 valid trials (i.e., 12 trials per digit/grapheme) were collected. Trials were followed by a 2 s interstimulus interval, indicated by a centrally presented ’x’ during which participants could blink or look away. In total, pupil responses of 5,760 trials were assessed in Block 1 in total, 1,920 from each of the three groups.

Only in synesthetes, we subsequently assessed pupil light responses to physically presented colors (Block 2, see Figure 1c). Block 2 was similar to Block 1, except that no digits were presented during the stimulus phase, but a centrally presented colored disc of 1.98 °visual angle in diameter, again on a dark gray screen. Ten different colors were presented in random sequence, one per trial. The color of the disc corresponded to the average per digit recreated color by the synesthete. Additionally, a smaller gray square was presented in the center of the disc, corresponding the same grayness value and number of pixels as the corresponding digit during Block 1. I.e., if a participant chose an on-average bright-blue for a ’5’ in Block 1, they were presented with a bright-blue in Block 2 with a central gray box corresponding to the grayness and number of pixels contained in the ’5’. Just as for Block 1, trials were followed by an interstimulus interval. Again, participants had to keep gaze in the center and not blink (trials with blinks were repeated in random sequence). Every ten trials, a pause screen allowed participants to take a break. Per color, 5 trials were assessed, i.e., 50 trials in total. In total, pupil responses of 800 trials were assessed from synesthetes in Block 2. The experiment took about 60 minutes in total for synesthetes and about 15 minutes for passive controls (40 for active controls).

### 4.4 Data processing

All data were processed using custom Python (v3.10) and R (v4.4.3) scripts.

#### 4.4.1 Pupil data

Pupillometric data were preprocessed following [48, 92]. Data were first filtered for valid trials only and down-sampled to 100 Hz. Pupillary data were transformed from arbitrary eyelink units to millimeters using a conversion factor obtained with an artificial eye [see 93]. Data were then subtractively baseline corrected using the mean pupil size during the last 50 ms of the baseline directly preceding the stimulus phase. Baseline pupil sizes did not differ between groups (*F*(2, 45) = 0.707, *p* = 0.499). Statistical tests over time were not corrected for type-1 error, which is why we recommend caution before interpreting short and briefly significant intervals strongly and to rely on analyses performed on averages per timebin in doubt.

#### 4.4.2 Color reports

Per-trial reported colors in RGB space were converted to HLS (hue, lightness, saturation) color space. The lightness domain was used to infer the qualia of synesthetic color. Color consistency was calculated following [17] under slight adjustments. For each participant and grapheme, the mean pairwise Euclidean distance between all twelve RGB color selections was computed and subsequently averaged. Smaller values indicate smaller color distance and thus higher internal color consistency.

#### 4.4.3 Questionnaire data

Questionnaire data (excluding the screening questionnaire) were collected on paper before the start of the eye-tracking task; responses were later digitalized. We calculated the projector-associator (PA) score following Rouw & Scholte [49]. Furthermore, we assessed the subjective strength of the coupling between each grapheme and color (referred to as ’coupling strength’ elsewhere; see Supplementary Materials for the questionnaire).

## Acknowledgments

We thank Tessie Hamers for assistance with piloting and all participants for their contribution.

## Data availability and code availability

Full materials, data, and analyses are available via the Open Science Framework https://osf.io/b6d8j/.

