## Supplementary materials for "Pupil size reveals the perceptual quality and effortless nature of synesthesia"

### Additional information on color reports

Supplementary Figure 1 visualizes hues and color reports to complement Figure 2 of the main manuscript. Note that colors appear visually more clustered for synesthetes and more often at full saturation (outer line) than for controls.

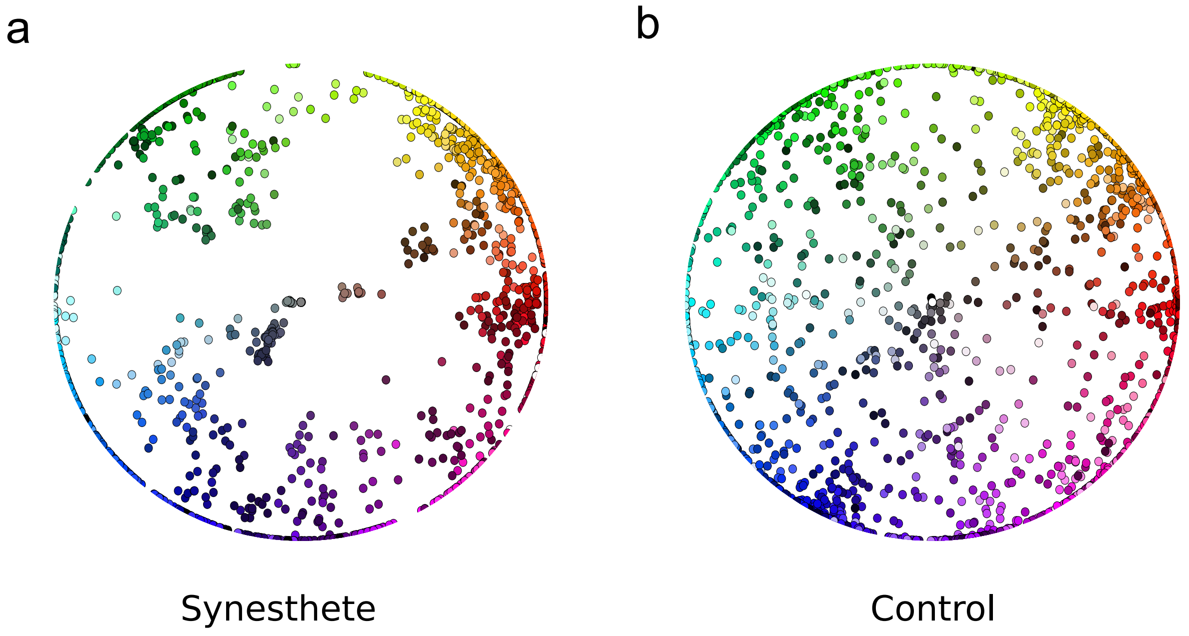


Supplementary Figure 1: Hue (angle) and saturation (eccentricity) for color reports for **a** synesthetes and **b** controls

### Pupil responses per grapheme

Supplementary Figure 2 gives pupil responses per grapheme and group of participants. No consistent patterns were found to graphemes across groups.

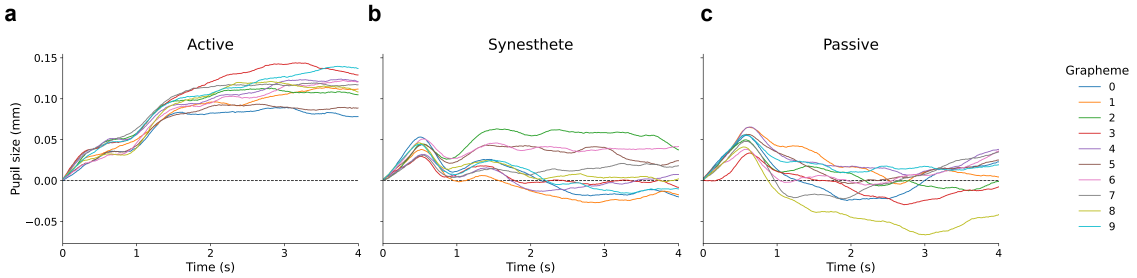


Supplementary Figure 2: Pupil size change to graphemes, irrespective of reported color. Horizontal dashed line represents baseline pupil size. **a** Active controls, **b** synesthetes, **c** passive controls.

### Pupil responses, lightness split without data exclusion

Supplementary Figure 3 gives pupil responses for the median split along the lightness dimension in Block 1 as in the main manuscript, but without any participants excluded for low trial numbers in either bin. The statistics support the same conclusions as for the arguably more sensible, but somewhat arbitrary exclusion presented in the main document.

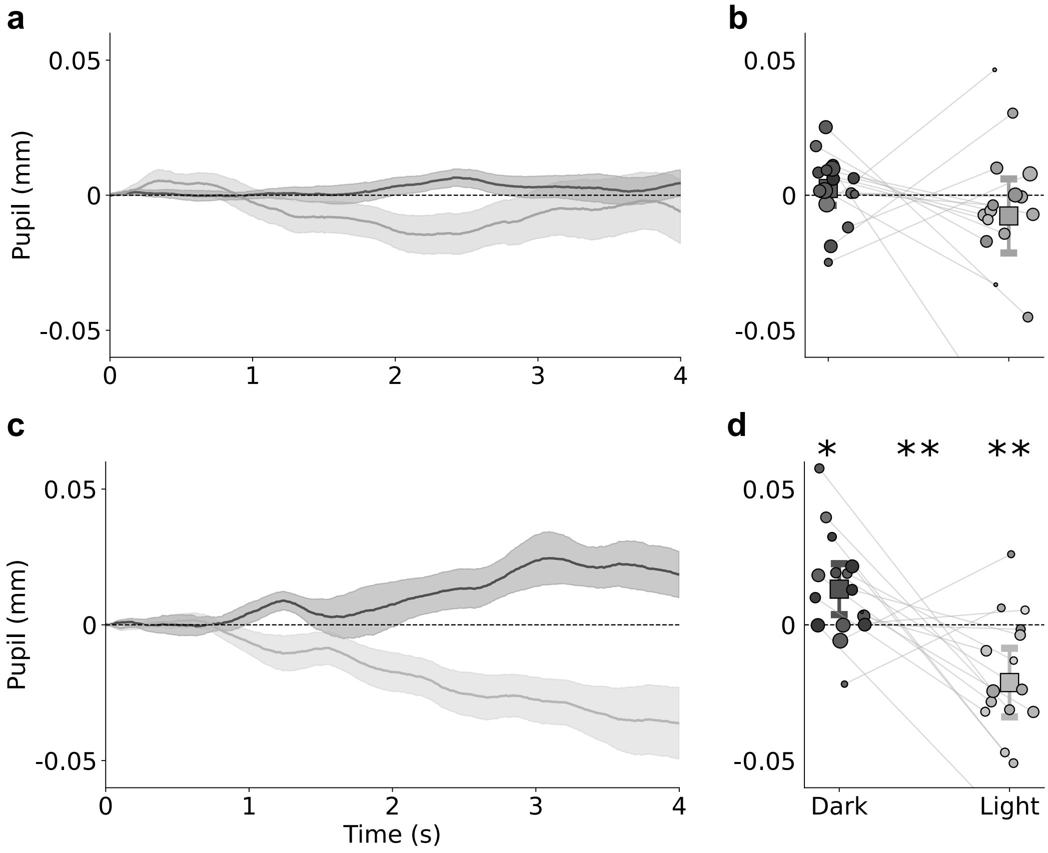


Supplementary Figure 3: Pupil size change to graphemes, split by 0.5 reported color lightness (dark gray = low lightness; light gray = high lightness) without removing participants with little trials per bin. Top row: pupil responses to graphemes in controls. Bottom row: pupil responses to graphemes in synesthetes. **a, c** depict average, baseline-corrected and within-participant demeaned pupil responses. Shaded error bands represent ±1 SEM across participant means. **b, d** depict mean pupil size (800–4000 ms) for dark vs. bright colors. Dots show individual participants; squares denote grand means with 95% CIs as whiskers. Dot luminance corresponds to the participant's average photism lightness per bin, dot size to the number of trials in the respective bin. *: $p < .05$, **: $p < .01$. Asterisks denote significance relative to 0 for lightness bins (left, right) and for the difference between lightness bins (center).

### Pupil responses without demeaning

Supplementary Figure 4 gives pupil responses for the median split along the lightness dimension in Block 1 as in the main manuscript, but without pupil size changes being demeaned to isolate effects of reported lightness.

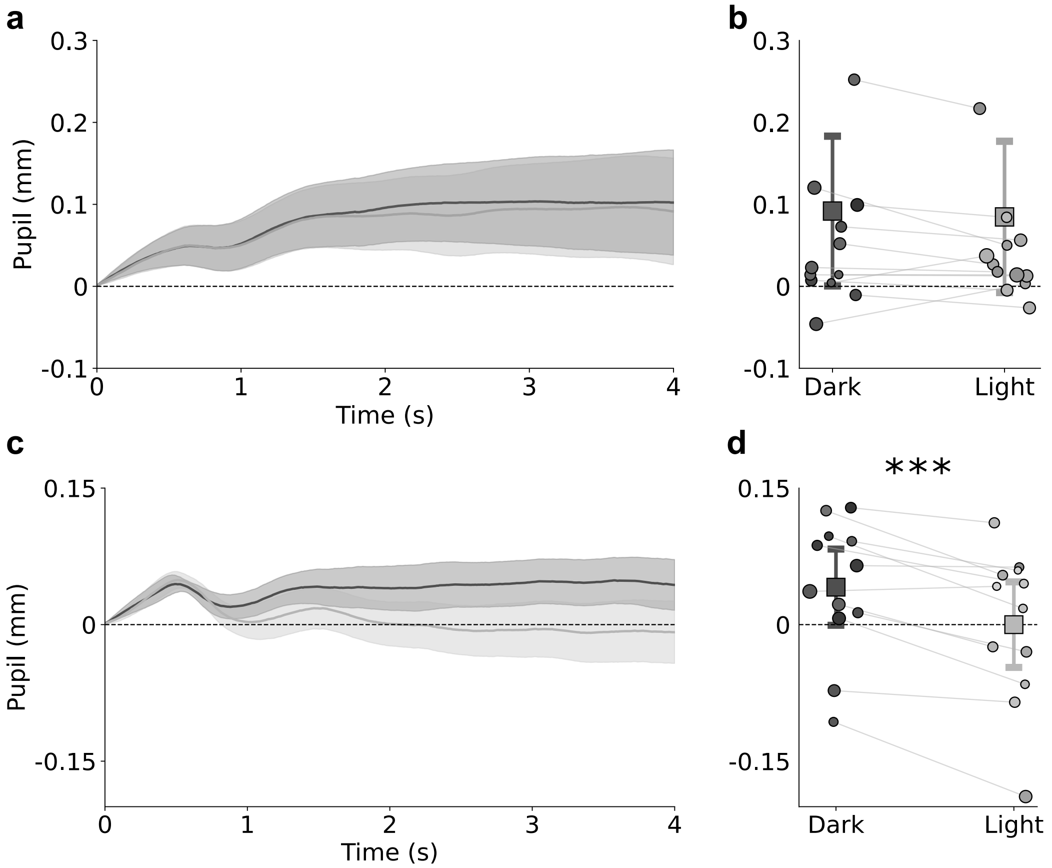


Supplementary Figure 4: Pupil size change to graphemes, split by 0.5 reported color lightness (dark gray = low lightness; light gray = high lightness) without demeaning (i.e., removing the average pupil response shape in the 4s stimulus interval per individual irrespective of brightness perception). Top row: pupil responses to graphemes in controls. Bottom row: pupil responses to graphemes in synesthetes. **a, c** depict average, baseline-corrected pupil responses. Shaded error bands represent ±1 SEM across participant means. **b, d** depict mean pupil size (800–4000 ms) for dark vs. bright colors. Dots show individual participants; squares denote grand means with 95% CIs as whiskers. Dot luminance corresponds to the participant's average photism lightness per bin, dot size to the number of trials in the respective bin. ***: $p < .001$. Asterisks denote significance between lightness bins (center).

### Lightness effects across coupling strength bins

Supplementary Figure 5 visualizes demeaned pupil responses to high/low lightnesses as obtained from the same split as applied in the main manuscript, but further split by reported coupling strength.

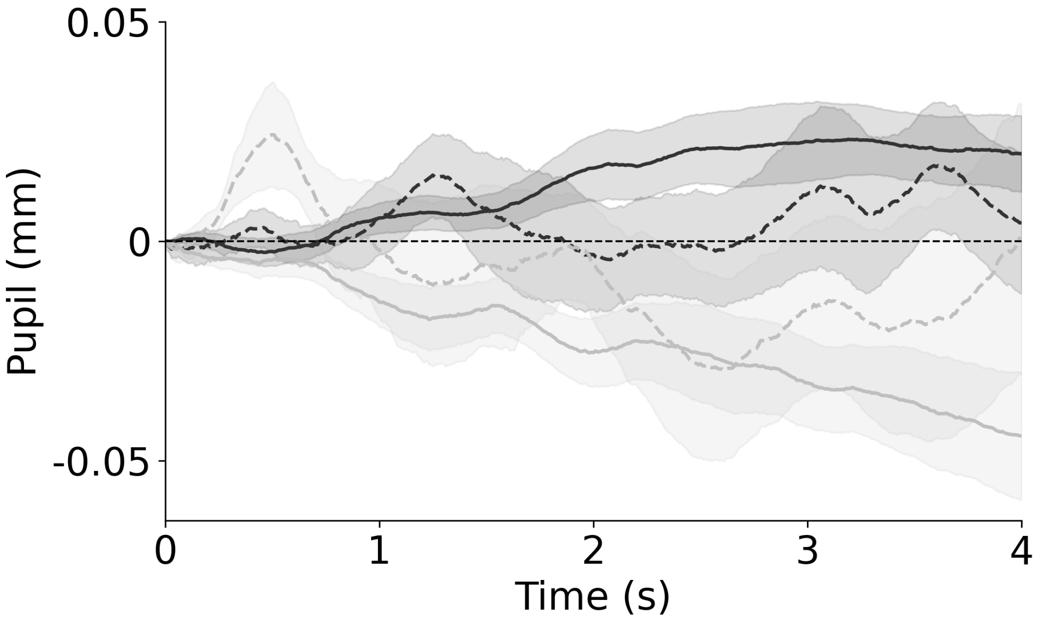


Supplementary Figure 5: Demeaned pupil responses to digits eliciting dark (black) versus bright (gray) lightnesses (0.5 cutoff). Solid lines depict responses for graphemes rated with high coupling strength (3,4, strong), dashed for graphemes with low coupling strength (1,2, weak). Participant counts per line: weak dark: n = 11, weak light n = 10, strong dark n = 15, strong light n = 15.

### Color consistency vs. coupling strength

Synesthetes: To test whether coupling strength and/or reported color consistency predict pupil size through lightness, a LME was run with coupling strength ($pupil_{800\text{–}4000ms}$ $\sim$ grapheme + lightness + coupling strength + color consistency + color consistency * lightness + lightness*coupling strength + (1 | participant). Color consistency did not, directly ($t$ = 1.522, $p$ = .128) or in interaction with lightness predict pupil size ($t$ = -0.670, $p$ = .503). As expected, coupling strength had no main effect on pupil size either ($t$ = 1.250, $p$ = .211). Crucially, however, both lightness ($t$ = -3.290, $p$ = .001) and the interaction of lightness with coupling strength affected pupil size ($t$ = -2.397, $p$ = .017). This implies that the consistency of reported colors, within synesthetes is not predictive of pupil size. Rather, the reported strength of synesthetic perception is, by increasing effects of lightness on pupil size. Or in other words: synesthetes see colors stronger that they report to see stronger, irrespective of how consistently they indicate them. Note, however, that variability in color consistency was very limited in synesthetes. Together, these analyses indicate that consistency of reported colors is insufficient to determine the strength of qualia in synesthetes.

In controls, using the same model as specified above for the synesthetes, neither lightness ($t = -0.947$, $p = .344$), coupling strength ($t = 0.427$, $p = .669$), color consistency ($t = 0.073$, $p = .942$), nor the interaction of coupling strength with lightness ($t = -1.033$, $p = .302$) or the interaction of color consistency with lightness ($t = -0.814$, $p = .415$) predicted pupil size. This demonstrates that reporting a high coupling strength or reporting a color consistently is not sufficient to produce synesthete-like pupil responses. This data speaks to the longstanding question as to whether synesthesia is 'just' a consistent association. In so far pupil responses reflect the visual perception described (as demonstrated for synesthetes), consistent associations are insufficient to evoke similar-to-synesthetic perception.

### Pupil responses to graphemes in active controls

Supplementary Figure 6 visualizes t-values of a per-timepoint LME for data of the active control group, specified as for synesthetes, but without the not-assessed PA score as predictors. We warn before interpreting the interaction reaching significance for a short period, given the absence of significance across the whole interval and short duration and significance level (akin to the interaction of PA score and lightness in synesthetes).

### Pupil response to physical luminance in synesthetes (Block 2, colored discs)

Supplementary Figure 7 provides the same per-timepoint linear mixed effects model as Figure 4 in the main manuscript, but for Block 2 (colored discs). As can be seen, the lightness of discs affected pupil size significantly as soon as 330 ms after onset. See Figure 3e of the main manuscript for a corresponding visualization of the pupil response, split by grapheme lightness.

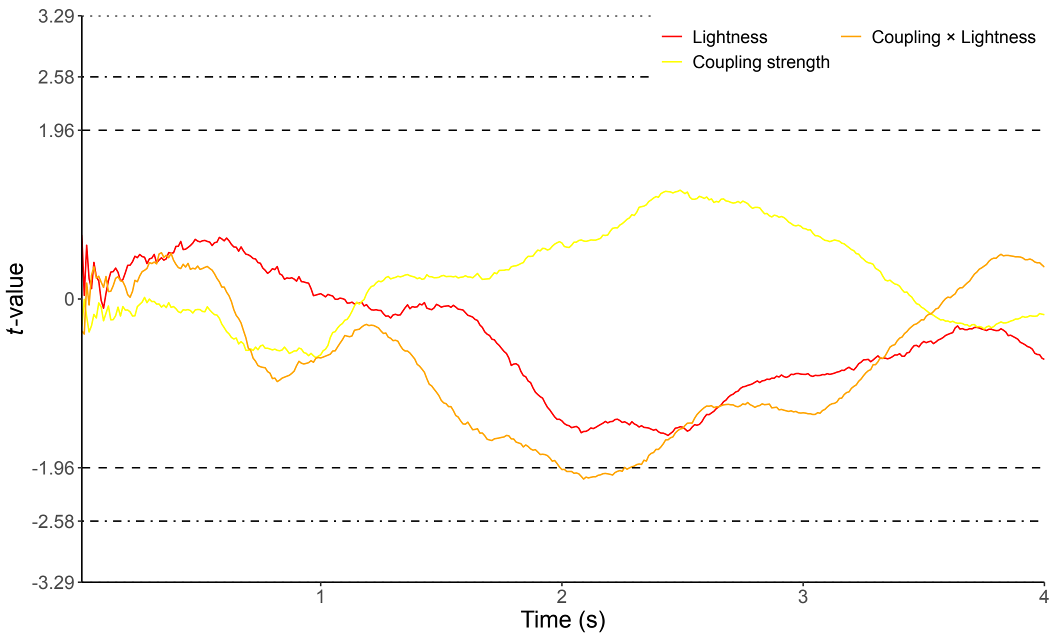


Supplementary Figure 6: per time point LME as Figure 4 in the main manuscript (intercept and graphemes not shown) for the control group indicating colors. Without PA score predictor, as PA score was not assessed in controls.

### Coupling strength questionnaire

Supplementary Figure 8 gives the questionnaire used to assess coupling strength per grapheme and synesthete.

### Screening questionnaire

Lastly, Supplementary Figure 9 shows the screening questionnaire used for recruitment/screening of synesthetes. Besides, we asked for contact information, name, and their willingness to travel to the laboratory to partake in the experiment.

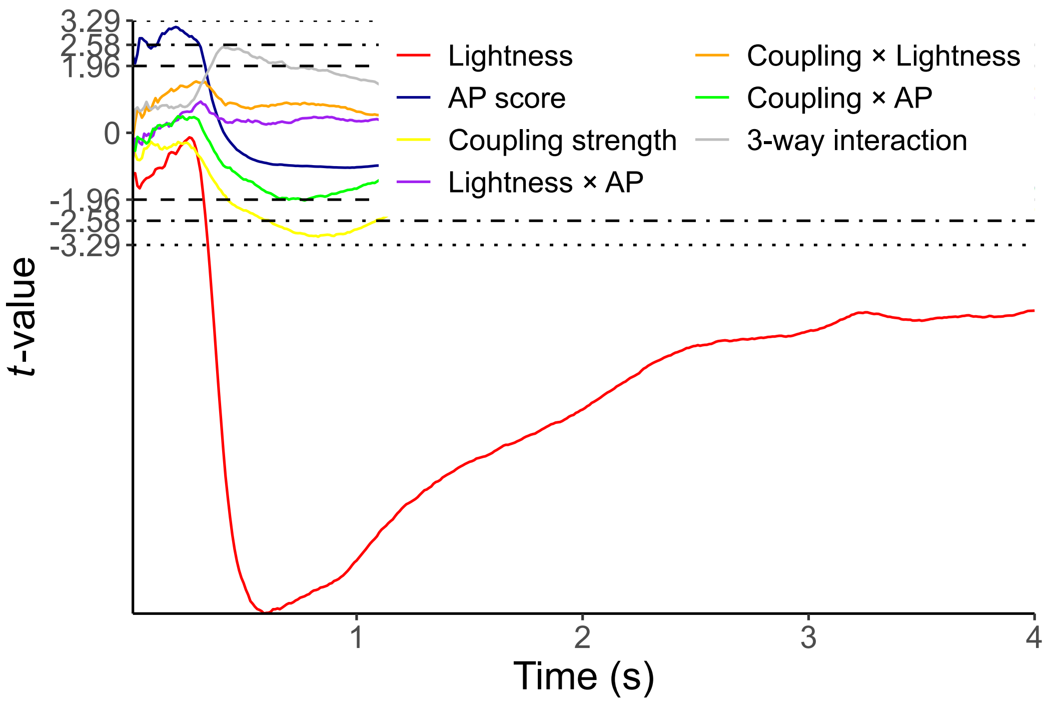


Supplementary Figure 7: Results of per-time-point linear mixed effects model predicting pupil size in synesthetes in Block 2 (colored discs). Covariates for the individual graphemes and intercept are not visualized here. All visualizations as in Figure 4 of the main manuscript.

**Please write down the numbers 0 to 9 in the table below to indicate how strong you associate a color to each individual number.**

| **Association/visualisation to color** | **Number** |
|---|---|
| **No** association/visualisation to color | |
| **Weak** association/visualisation to color | |
| **Moderate** association/visualisation to color | |
| **Strong** association/visualisation to color | |
| **Very strong** association/visualisation to color | |

Supplementary Figure 8: Questionnaire used to assess the coupling strength between grapheme and color in synesthetes and active controls.

### Your Experience With Synesthesia

For my research, I'm looking for **people in The Netherlands, that experience Grapheme-Color Synesthesia**.

This form is meant to get an overview of possible candidates to participate in an experiment in which I attempt to measure the strength of synesthetic experiences, and create the first physiological measurement method for synesthesia. By filling in this form, you are not bound to anything.
**If you experience Grapheme-Color Synesthesia, and you live in The Netherlands, it would help me enormously if you could fill in this form**. If you know others that experience synesthesia, please send them this form too.

Filling in this form will not take longer than 5 minutes

If you have any questions, you can contact me at

**Do you experience Grapheme-Color Synesthesia?**
*Grapheme–Color Synesthesia is a form of synesthesia in which an individual's perception of numbers or letters is associated with the experience of colors.*

- ○ Yes
- ○ No
- ○ I'm not sure

The following questions are about your experiences with Grapheme-Color Synesthesia.

**Do you experience colors with numbers, letters, or both?** *

- ○ I experience colors with numbers only.
- ○ I experience colors with letters only.
- ○ I experience colors with both numbers and letters.

**How consistent are your synesthetic experiences?**

| | 1 | 2 | 3 | 4 | 5 | 6 | 7 | 8 | 9 | 10 | |
|---|---|---|---|---|---|---|---|---|---|---|---|
| Very inconsistent. Example: the number 5 always results in experiencing a different color. | ○ | ○ | ○ | ○ | ○ | ○ | ○ | ○ | ○ | ○ | Very consistent. Example: the number 5 always results in experiencing in the same color |

**Which statement applies the most to you?**

When reading numbers or letters...

- ○ ...I see a color projected onto the number or letter itself.
- ○ ...I see a color somewhere else in the space around me.
- ○ ...I 'see' or picture a color in my mind's eye.
- ○ ...I associate these numbers or letters to a color, but I do not see or picture this color.

Supplementary Figure 9: Screening questionnaire to identify and recruit participants with grapheme-color synesthesia.
